# A Patient-based Model to Investigate the Effect of Micro-structure Scars on the Optogenetic Defibrillation Success: A Computational Modelling Approach

**DOI:** 10.1101/2020.08.13.249250

**Authors:** Elham Zakeri Zafarghandi, Fariba Bahrami

## Abstract

This study investigates the role of the microstructure of real scars in the success of optogenetic defibrillation. To reduce the computational cost of high-order models (like Ten Tusscher Model, TTM) for a single cell as well as to take advantage of their ability to generate a more realistic output, we developed a low-order model of optogenetic cardiac tissue based on the modified Alieve-Panfilov single-cell model and estimated its parameters using a TTM. Two-dimensional electrophysiological cardiac tissue models were produced including different scar shapes that were extracted from Late Gadolinium-Enhanced (LGE) magnetic resonance imaging data set of 10 patients with non-ischemic dilated cardiomyopathy. The scar shapes were classified based on four criteria: transmurality, relative area, scar entropy, and interface length. Scar with the highest 25% of the relative area showed 25% of successful cases, this ratio is 27%, and 25% for a scar with the most top 25% of entropy, and transmurality, respectively. In comparison, the proportions are 61.54%, 44.44%, and 61.76%, for the lowest 25% of the area, entropy, and transmurality. We also investigated the efficacy of various methods for light-sensitive cells’ distribution within the cardiac tissue with scar. Four types of distributions were defined. Defibrillation within tissues with 0.1 light-sensitive out of all cells was 15 to 25% more successful than their counterparts with 0.05 light-sensitive cells. Lastly, we examined the effect of an earlier stimulation on the success probability of defibrillation. Our results indicated that inducing 0.5 msec earlier resulted in a roughly 15% rise in successful cases.

## 1. Introduction

Ventricular fibrillation is one of the most prevalent heart arrhythmias, which might result in blood clots, stroke, or even in serious conditions, heart failure. As a conventional treatment, an implantable cardioverter-defibrillator (ICD) is used for a heart prone to arrhythmia. Unfortunately, ICDs have some undesirable side effects, including structural damage to cardiac tissue, which, in turn, would result in a more heterogenous cardiac tissue. These side effects have inclined scientists to find new medical approaches addressing fibrillation problems. Optogenetic treatments (defibrillators) are convenient candidates to substitute for cardiac therapies based on electrical stimulations of the cardiac tissue [1, 2].

In 2005, Li Li *et al*. examined the mechanism of shock-induced arrhythmogenesis and arrhythmia maintenance in a rabbit model of a healed myocardial infarction. They proved that an infarcted rabbit heart is more vulnerable to shock-induced arrhythmogenesis and arrhythmia maintenance [3]. Also, in another research [4], utilizing MRI and DTI data, researchers developed a 3-D model of the infarcted canine heart, by which they claimed that borders between intact and scar tissues are the location from which subsequent fibrillations will start, because of their resulting heterogeneity. Regarding the fact that a considerable number of candidates for ICD implantation are who have a prior myocardial infarction, the evaluation of the efficacy of treatments in such conditions would be crucial. In [5], a 3-D model was developed, and then the authors showed that electrical defibrillators have lower chances of being successful in infarcted hearts. According to the difference between the electrical and optical stimulations, understanding the effect of optical defibrillating shock within an infarcted cardiac tissue is highly essential.

In this study, we developed a low-order computational model using LGE_MRI data of non-ischemic dilated cardiomyopathy patients. Utilizing this model, we sought to investigate the efficacy of optical defibrillation, while investigating the effect of the fibrosis microstructure and also the distribution of light-sensitive cells through cardiac tissue. First, we simulated the action potential of normal and optogenetic single cardiac cells using the Ten Tusscher cardiac cell model (TTM). This data, because of the precise output of the TTM, formed our main data set. Using this data set, we estimated the parameters of a modified Aliev-Panfilov (AP) single cardiac cell model. Afterward, having the optogenetic cells spread stochastically among normal cells, we developed the model of the optogenetic ventricular tissue without scar. In this step, in order to validate our proposed model, we compare the features of spiral waves, as an indicator of fibrillation, between our model and the TTM. After the validation step, we inserted scar tissue extracting from LGE-MRI data into our model. Lastly, we induced fibrillation through the tissue and evaluated the efficacy of optical defibrillation under the condition of different types of scars extracted from the data set [6].

## 2. Materials and Methods

### 2.1. MRI Data Set

In this study, we have utilized the LGE-MRI data of 10 patients with non-ischemic dilated cardiomyopathy [6] with the scanning resolution of 0.64-1.97mm in-plane, and 7-10.5 mm slices thickness. Fig 1 shows one of the LGE-MRI accompanied by extracted scar shape from that image.

**Fig 1.**
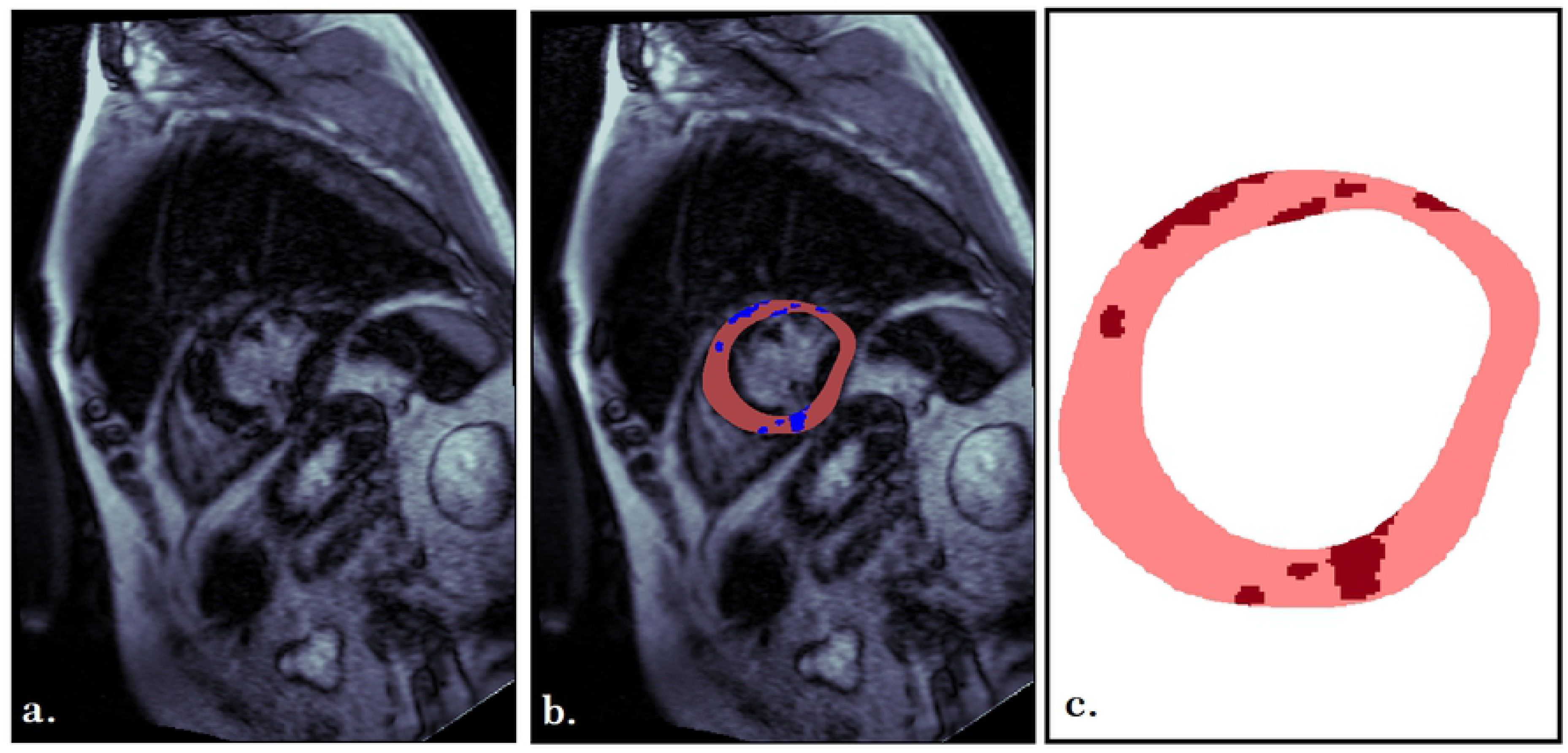
Scar’s shape extracting from LGE-MRI data. a. LGE-MRI raw data b. scar’s shape extracted from “a” (blue pixels are indicators of fibrosis, and pink marked pixels are the cardiac cells) c. consequent scar’s shape (red pixels are indicators of fibrosis, and pink pixels are the cardiac cells).

### 2.2. Scar shape

We have used 2-D patterns of LGE extracted from 47 LGE-MRI images of 10 patients. Four features of each extracted scar shape have been measured, including transmurality, relative scar area, LGE entropy, and interface length of LGE-myocardial. We have selected these features because of the significant roles they play on the electrical behavior of the cardiac tissue. Transmurality may well manipulate the chance of reentry phenomenon, relative area could be important for tissue conductivity, and entropy and interface length are important features for source-sink mismatch.

The transmurality was measured by the ray-tracing approach. The relative area of scar was defined as the ratio of enhanced to the non-enhanced myocardial pixels. The scar entropy was measured by Shannon’s standard entropy. And finally, the interface length was the sum of borders between myocardial, LGE, and borders arclength [6].

Table S1 in Supplementary section shows the average and interquartile amounts of the data set.

### 2.3. Computational modeling

#### 2.3.1. Channel Rhodopsin2 (Chr2) model

The model for Chr2, which we utilized in this study, is a four-state model introduced in [7]; the reason for this selection will be discussed later. Using four different states, the four-state model describes the dynamics of the Chr2; these states are as follows.

Closed sensitized (C1), Excited(O1), Open(O2), and Closed desensitized(C2).

#### 2.3.2. Optogenetic cell model

The Gene Delivery (GD) approach to make cells optogenetic does not affect normal electrical activities of a cell [8]; therefore, in this study, we assumed that the GD method was used in order to making cells optogenetic. We needed a set of data from two different types of cells, namely light-sensitive and typical cells. Because of utilizing the GD method, the light-sensitive cells have one light-gated channel more than normal cells. In this regard, we used the Ten Tusscher cardiac cell model [9] and added up the Chr2 model to yet other membrane’s currents to produce the model of a light-sensitive cell.

#### 2.3.3. Single-cell model

The Ten Tusscher cardiac cell model can mimic precisely the timing of a real cardiac cell. Although, because of the number of variables and dynamics, it leads to us encountering intricate computational problems for tissue modeling. Computationally speaking, in comparison with the Ten Tusscher model, the Aliev-Panfilov is a simpler model that can be used to model the cardiac tissue. Also, it is precise enough for describing cardiac fibrillation and defibrillation procedure, since in such a phenomenon, the exact shape of the action potential is not needed, and just passing the activation threshold is the issue. Therefore, in order to take advantage of the Ten Teusscher model, using the output of the resulting Ten Tusscher action potential for both light-sensitive and normal cells, we estimated parameters of a low-order cardiac model, the Aliev-Panfilov, in a way that it approximates the timing of Ten Tusscher action potential as well as possible.

The Aliev-Panfilov model is a simple model that consists of two dimensionless variables u and v describing fast and slow processes (1,2):

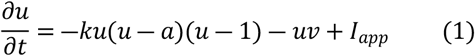

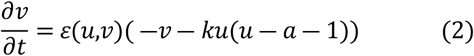

Where,

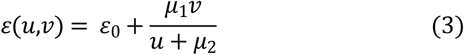

Where Iapp = 0.4 ^PA^/_PF_, *ε*0=0.002, k = 8, a = 0.15, *μ*1=0.2, and *μ*2=0.3 [10].

As discussed in [11], the parameter “a” is called the threshold; the parameter “*ε*” describes the ratio of the time scales of variables *u* and *v*.

The Aliev-Panfilov model, because of the *uv* term, which shapes the nullclines geometry, is more appropriate for describing the cardiac cell dynamics than the Fitzhugh-Nagumo model. In this study, we changed (2) to (4) [12]

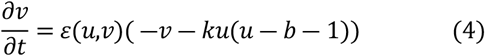

Where b = 0.05, *i.e*. the parameter “a” in (1) and (2) is different now. And the membrane voltage calculated as follows [12].

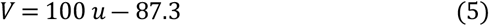

Fig 2 shows four different action potentials, namely resultant action potential of the Ten Tusscher model without Chr2, with the 3-state model of Chr2, and with the 4-state model of Chr2, and also the consequent action potential of Aliev-Panfilov model without mentioned modifications. As clearly can be seen in this figure, there is no principal difference between the final action potential of the cell after inserting the 3-state and 4-state model of Chr2. On the other hand, the 4-state model establishes a better insight into the photo-kinetics of ChR2-expressing cells [7]; therefore, we used the 4-state model in our *in-silico* simulations, which can keep possible the chance of further future modifications.

**Fig 2.**
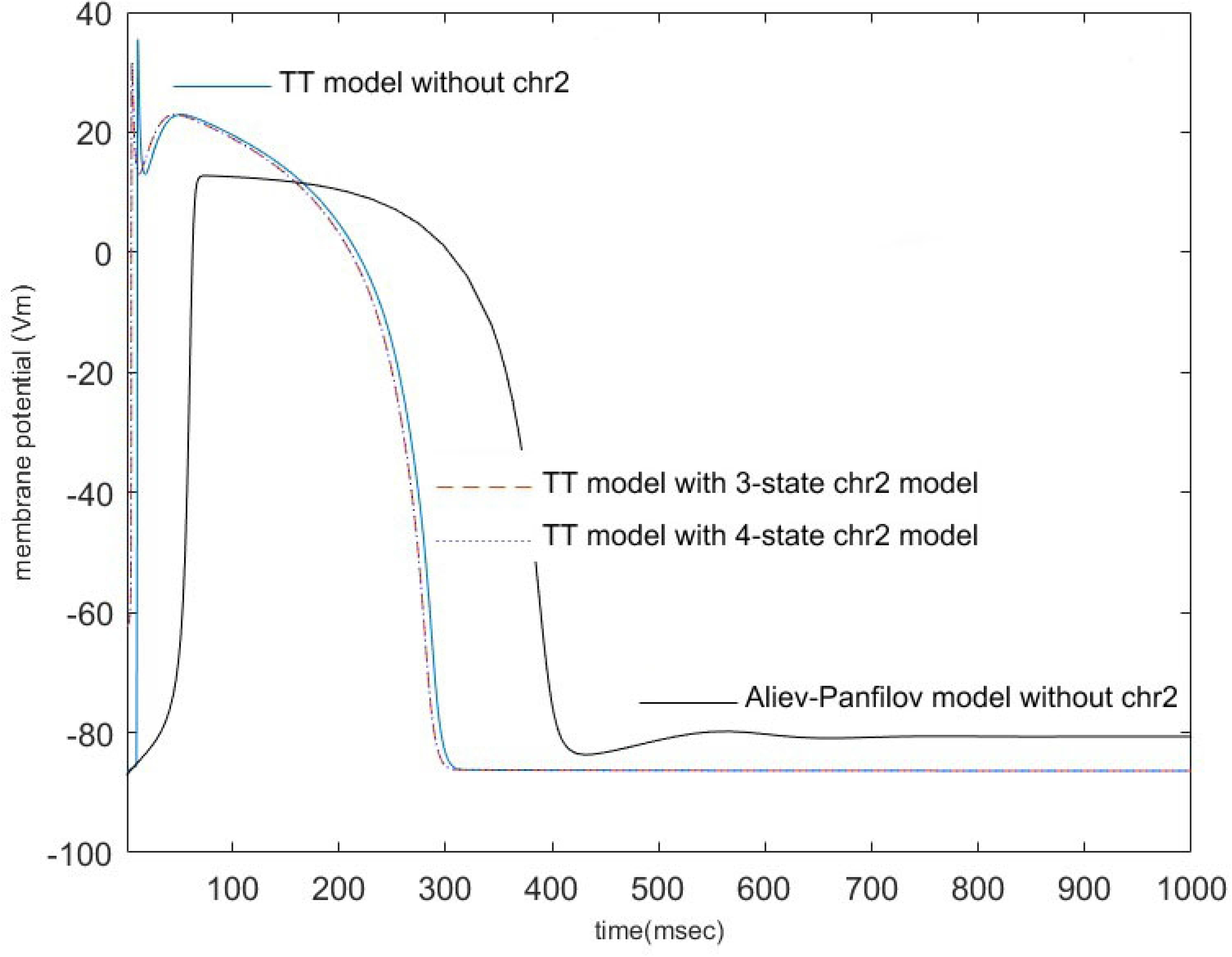
Resultant action potential of different models-solid blue line: TT model without Chr2, dashed red line: TT model with the 3-state model of Chr2, dot blue line: TT model with the 4-state model of Chr2, solid black line: Ap model without Chr2 before parameter estimation

#### 2.3.4. Parameter estimation

We used the genetic algorithm (GA) contained in MATLAB (R2019a) Optimization Toolbox to estimate the best set of parameters, which result in the lowest amount of error function. Summation of the squared differences was assumed as the fitness function with six variables (*i.e.*, k, a, *ε*_0_, *μ*_1_, *μ*_2_, and b). The initial value for parameters was as follows.

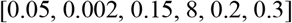

Table 1 shows the estimated parameters for normal and light-sensitive ventricular cells. And Fig 3 shows the final action potentials with estimated parameters.

**Table 1.** Estimated parameters of the Aliev-Panfilov Model

|  | **a** | **$\epsilon_0$** | **b** | **k** | **$\mu_1$** | **$\mu_2$** |
| --- | --- | --- | --- | --- | --- | --- |
| Normal Cell<br>(without Chr2) | 0.1 | 0.0043 | 0.08939 | 7.0717 | 0.011 | 0.3979 |
| Light-sensitive Cell (with 4-state model) | 0.1 | 0.0023 | 0.31 | 8 | 0.0011 | 0.7351 |

**Fig 3.**
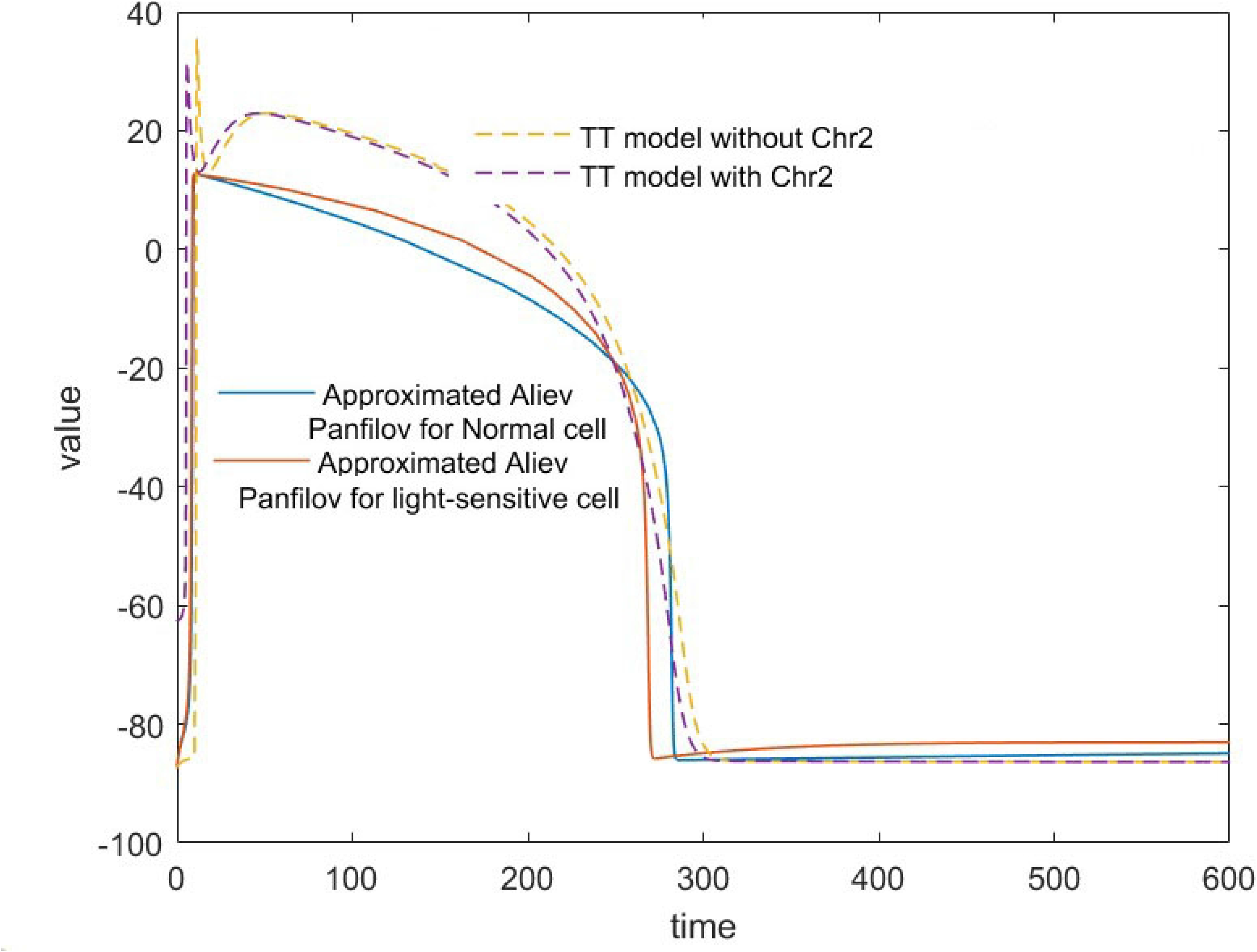
Resulting action potential of AP model with estimated parameters shown in Table1, compared with that of the TT model. It can be observed that, compared to AP action potential shown in Fig. 2, that of Fig. 3 is much more similar to the TT model’s AP.

#### 2.3.5. Tissue model

In the GD method, after systemic injection of modified cells in the laboratory, each cell will locate in a position in the target organ, stochastically. For simulating this process, we used the algorithm, which was introduced in [13] for fibrosis distribution for the first time, and after that was adapted for optogenetic approaches in another paper [14]. This algorithm consists of two main parameters P_thr_ and D, the indicator of patchiness and distribution, respectively. Fig 4 presents the distribution of light-sensitive cells within a 128*128 cell tissue for two different values.

**Fig 4.**
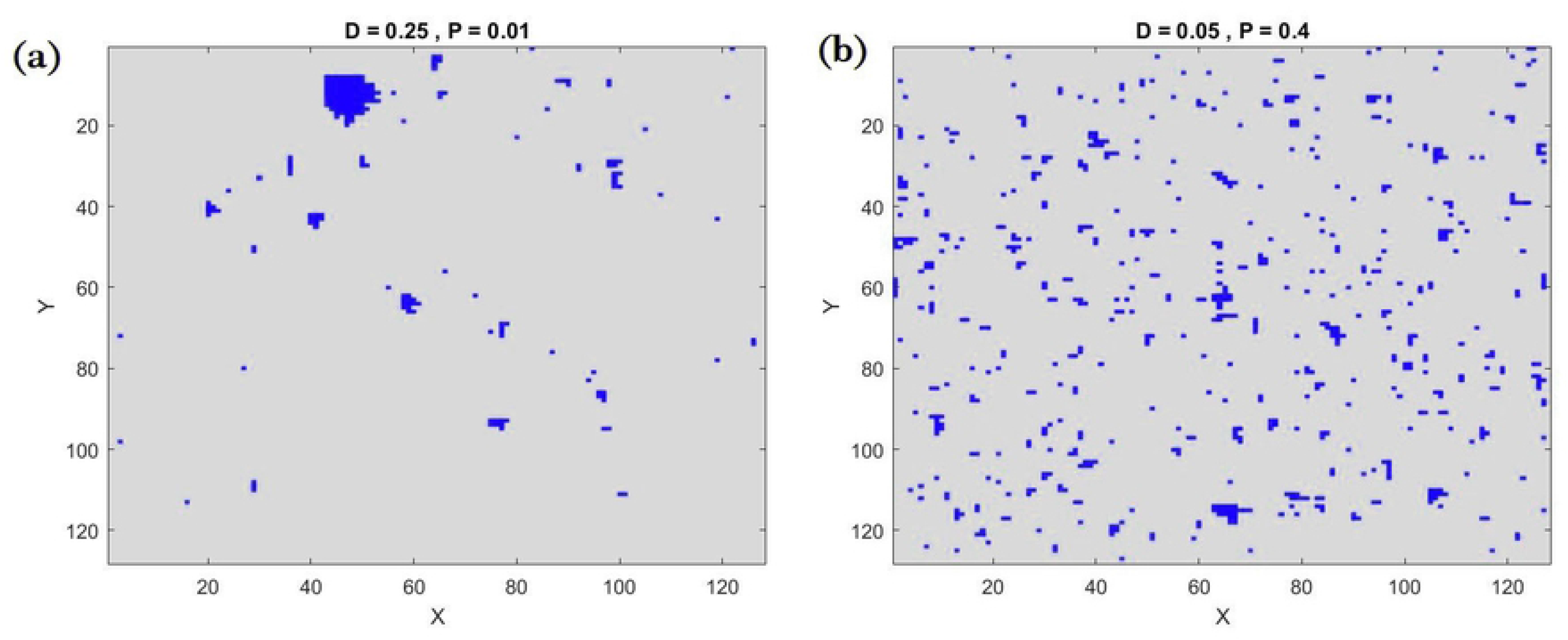
The distribution of light-sensitive cells within a 128*128 cell tissue_blue pixels are light-sensitive cells (a: D = 0.025 and P = 0.01, b: D = 0.05 and P = 0.4)

We sought to simulate the defibrillation procedure, so solving the bi-domain process was a crucial part of the study; since it can get us access to intracellular and extracellular spaces, independently. The bidomain problem equations are as follows.

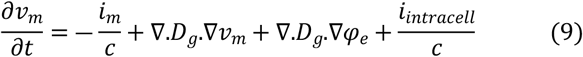

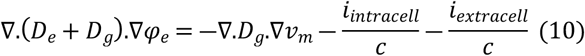

We utilized the five-stencil finite difference approach to solving these equations. Finally, the resultant equations are as follows.

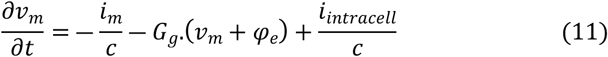

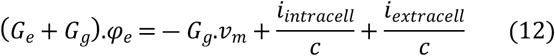

Where *φ*_*e*_ is the extracellular node voltage, *υ*_*m*_ is the membrane voltage (*φ*_*i*_ − *φ*_*e*_), *G*_*e*_ and *G*_*g*_ are intracellular and extracellular admitance matrix, respectively. C is the membrane capacitance and also *i*_*intracell*_, *i*_extracellular_ are injected currents into an intracellular and extracellular node. *i*_*m*_ is the current of membrane’s ion channels.

#### 2.3.6. Defibrillation

For defibrillation procedure, given the current which light stimulation induces in a single light-sensitive cell [15], we assumed an pproximately stable ionic current for each stimulation induction, *i.e.*, considering experimental data in [15] a pulse shaped light stimulation for 500msec with amplitude around 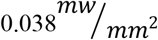 induce a current with a maximum of 12.01 ± 2.54 ^PA^/_PF_. In this study, we considered a 12 ^PA^/_PF_ current for each light-sensitive cell once the stimulation pulse is applied.

### 2.4. Code Availability

The codes for the computational model are publicly available upon request.

## 3. Results

### 3.1. Spiral wave’s tip movements

The movement of the tip of the spiral wave in real cardiac spiral waves have a part of linear shape [16,17], but simple models such as Fitz Hugh-Nagumo because of their rigid rotation are not able to mimic those movements.

We simulated the fibrillation process within a squared shape tissue for 30sec to compare the resultant spiral wave of our modelwith that of the tissue described by the Ten Tusscher single-cell model. In this regard, two different features of the spiral waves were considered, the average period of the spiral wave rotation, and the shape of its vortex core. The former showed the pace of wave propagation and the later presented the trajectory of that. As reported in [16], the average period of rotating for the resultant spiral wave within endocardium tissue is 264.23±10.37 msec, this value for the proposed tissue model (after averaging 60 time periods of spiral wave rotations) is 253.44±7.81 msec. Under this categorization, trajectory of the spiral wave tip of the proposed model is considered as quasilinear, which is similar to the results obtained by the Ten Tusscher model in [16]. Fig 5 shows the linear movement of the spiral wave’s tip from t=2sec to t=29.6180sec, *i.e.*, from timestep 20000 to timestep 296180 (the white line shows the trajectory of movements).

**Fig 5.**
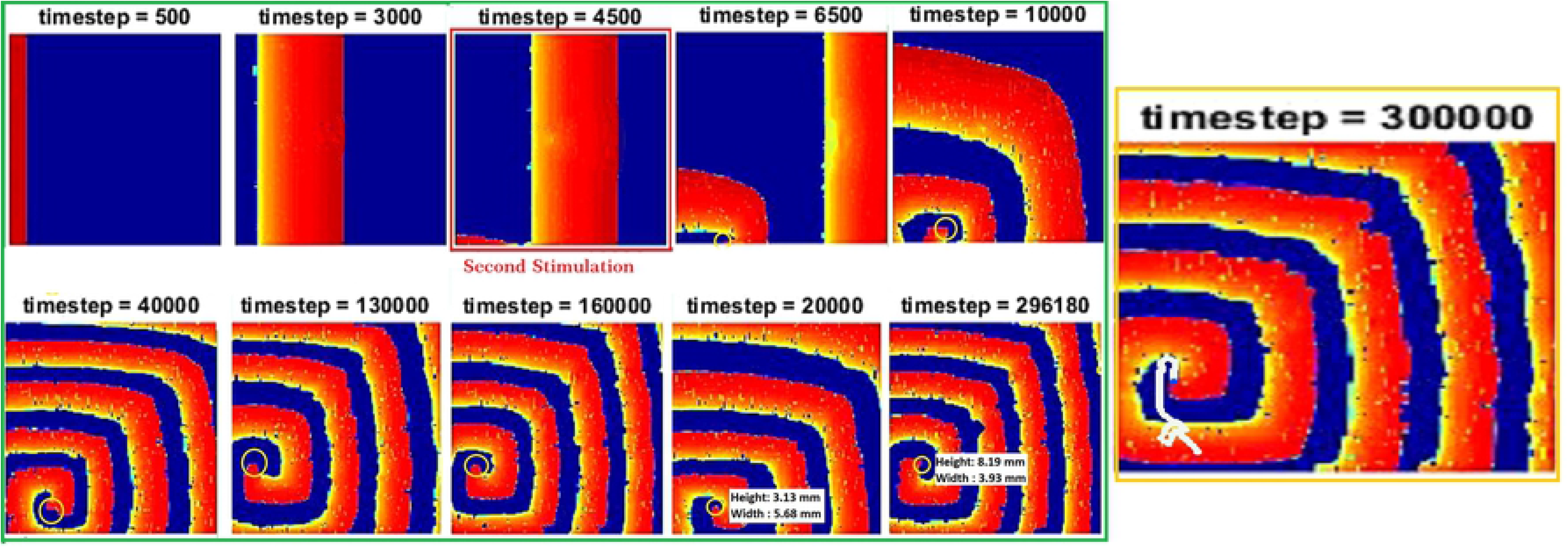
Green panel: The spiral wave formation in cardiac tissue with light-sensitive cells _ Orange panel: Trajectory of the tip of the spiral wave

### 3.2. Simulation of fibrillation

We induced fibrillation within each of the 47 models, extracted from LGE-MRI photos, under4 different conditions of light-sensitive cells’ distribution. First, low density and low patchiness (DLPL - D = 0.05 and P = 0.4), second, low density and high patchiness(DLPH - D = 0.05 and P=0.99), third, high density and low patchiness(DHPL - D = 0.1 and P = 0.4), forth, high density and high patchiness (DHPH – D = 0.1 and P = 0.99 ). Fig 6 Shows these four conditions for one of the models.

**Fig 6.**
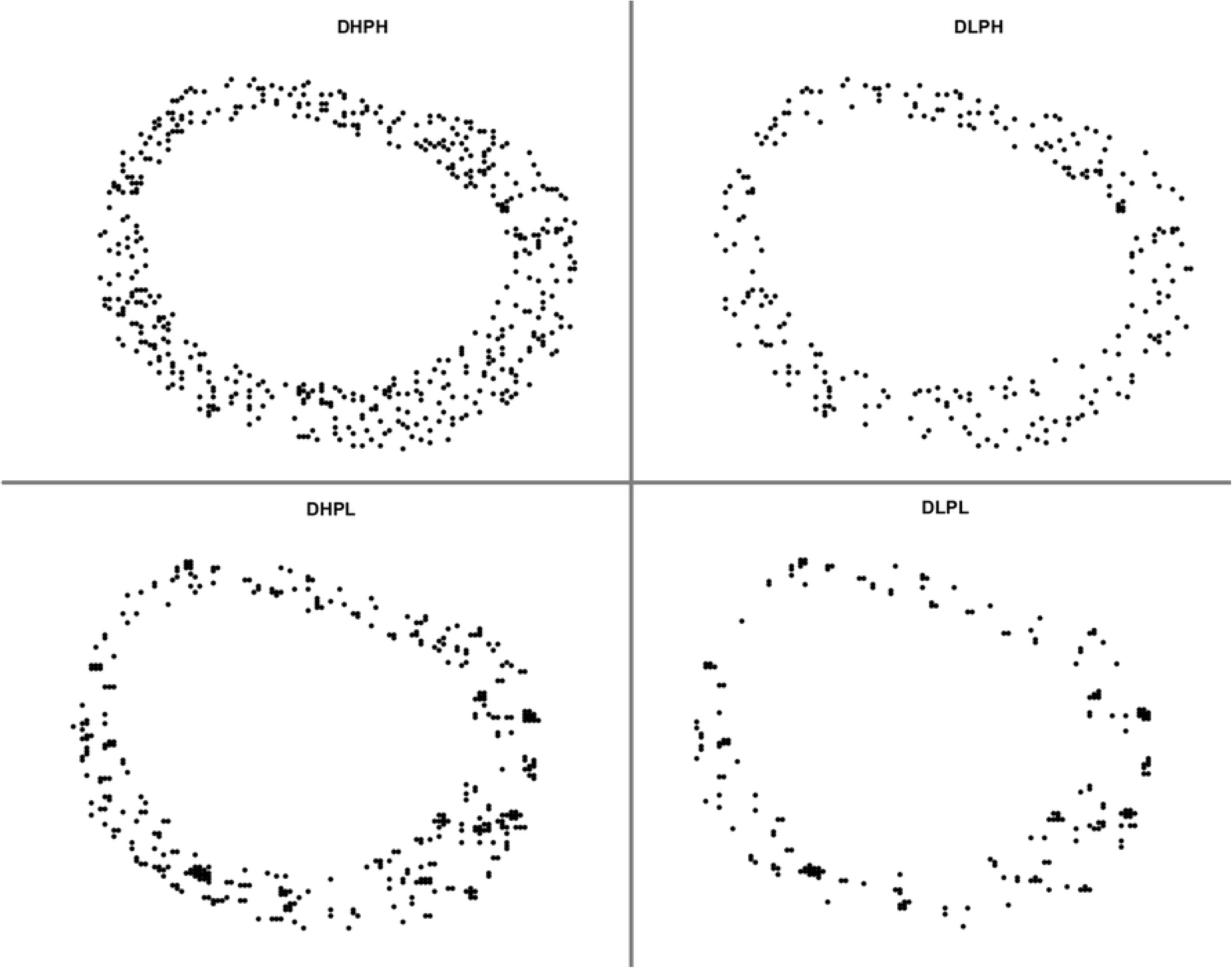
Distribution of light-sensitive cells with in the cardiac tissue – black dots are indicator of light-sensitive cell DLPL(D = 0.05 and P = 0.4), DLPH(D = 0.05 and P=0.99), DHPL(D = 0.1 and P = 0.4), DHPH(D = 0.1 and P = 0.99)

Each tissue was simulated for 30,000 time steps to be sure that the fibrillation is in stable circumstances and will not be damped after some time steps. The video of 4 cases out of 188 simulated ones is attached to supplementary materials, the Fibrillation folder. As illustrated in Fig 7, once the stimulation pulse produces, two wavefronts create and start traveling from the stimulation point trough cardiac tissue. So the cells in the pathway would be entered their inactivation period. When the second stimulation induces, because of fibrosis existence and the intrinsic low conductivity of fibrosis tissue in comparison with healthy tissue, it is likely to result in zones of transient conduction block, which restricted traveling waves into narrow paths. Once the wavefront from these narrow paths encounters a wide inactivated area, the source-sink mismatch may well stop the electrical wave propagation.

**Fig 7.**
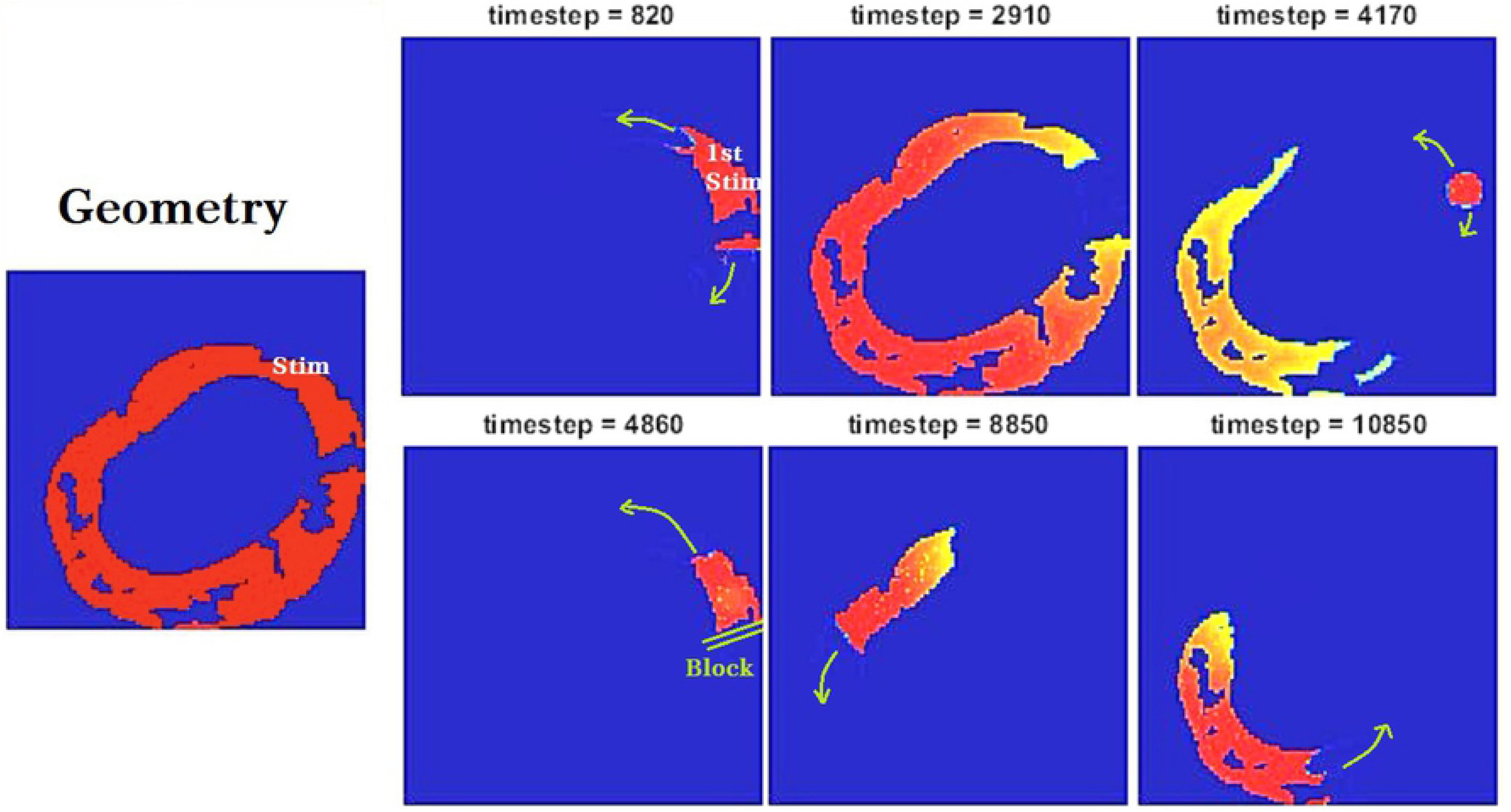
fibrillation simulation steps-Left panel: Geometry of the cardiac and fibrosis tissue, and stimulation location

### 3.3. Simulation of defibrillation

For the defibrillation procedure, we injected the mentioned current to the light-sensitive cells within the cardiac tissue at two different time steps, which is, first, step 20,000 (2 msec after the first stimulation) for all four categories, namely DHPH, DLPH, DHPL, and DLPL, and second, step number 15000 (1.5 msec after the first stimulation) for the DLPL category (called DLPL_qiuckstim in this study).

We defined successful defibrillation as a stimulation which scilences (damps) entirely the spiral waves. Based on this definition, the percentage of successful defibrillations among all cases was calculated and shown in Table 2. As clearly can be seen, the DLPH distribution of light-sensitive cells has a considerably lower amount of fibrillations being completely silenced.

**Table 2.** Percentage of successful defibrillations

| DHPH | DHPL | DLPH | DLPL | DLPL_quickstim |
| --- | --- | --- | --- | --- |
| 50 | 57.14 | 25 | 42.85 | 57.14 |

### 3.4. Defibrillation within tissues with different types of scars

As discussed in Materials and methods, for each scar shape extracting from LGE-MRI data, we calculated the amount of four features. Also, we divided each feature into four quartiles, namely 1^st^, 2^nd^, 3^rd^, and 4^th^ quartile. Table 3 shows the percentage of successful optogenetic defibrillation within each of the mentioned categories. The video of 5 cases out of 235 simulated ones is attached to supplementary materials, the Defibrillation folder.

**Table3.**
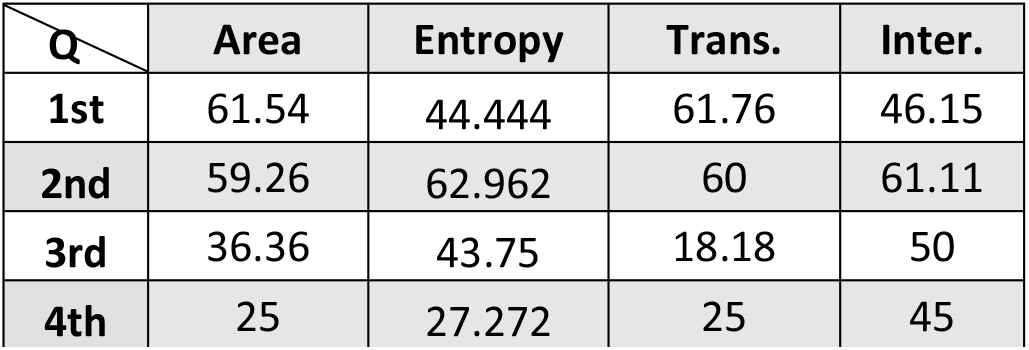
Percentage of successful defibrillation regarding each quartile of scar’s features

## 4. Discussion

### 4.1. The effect of scar’s micro-structure on the defibrillation success

We sought to investigate the efficacy of optogenetic defibrillators in more realistic circumstances. So we investigate the issue in the context of non-ischemic dilated cases. We categorized scars extracting from LGE-MRI images via four criteria, namely relative area, entropy, transmurality, and interface length.

As far as “Relative Area” is concerning there is a considerable change between the percentage of successful defibrillation cases within the first half and the second half of our data set, *i.e.* the probability of successful defibrillations is about 60% in the case of the area of the scar being less than 0.06 of the tissue, while this percentage would be about 30% for the scar area more than that. This issue could be discussed through the lower conductivity of fibrosis tissue. As discussed in [18, 19], two ex-vivo studies, the non-ischemic fibrosis, result in a decline in the velocity of conductivity. Therefore, once the stimulation produced by optic induction of light-sensitive cells enter a fibrosis area, it slows down there and when it could exit that area all neighbors are in their active period so the reentry phenomenon could occur. It means that, although the tip of the previous wave would be eradicated entirely, newly-generated wavefronts would be formed.

Regarding entropy, there is a significant drop in the chance of successful defibrillation in the 4^th^ quartile. Literally, the greater entropy, the higher variation exists in the microstructure of scar; so the source-sink mismatches would be deteriorated, and the abovementioned phenomenon results in forming new wavefronts. In our study, there is a rise from the 1^st^ to 2^nd^ quartile; we think that is because of the small amount of difference in the three firsts quartiles of our data set.

These results are in agreement with [20], as Muthalaly *et al.* concluded that high LV entropy is a predictor of forthcoming arrhythmias.

As discussed in [21], the less transmurality in the micro-structure of scar tissue may well reduce the chance of reentry phenomenon; the transmurality can lessen the probability of occurrence of successful defibrillation. The point at issue is that the less transmural scars within a tissue are equal to the less internal activation paths, which could, in turn, reduce the consequent reentry after defibrillation stimulation.

We could not see any meaningful relation between interface length and the percentage of successful defibrillation. Although in previous studies, the role of the scar interface was discussed.

### 4.2. Efficient structure for light-sensitive cells’ distribution

We sought to suggest an efficient structure for light-sensitive cells being distributed within the cardiac tissue. In this regard, we examined four different structures. As clearly can be seen in Table2, the defibrillation within tissues with higher amounts of light-sensitive cells (for D = 0.1) was about 15% to 25% more successful than their counterparts with lower amounts of light-sensitive cells. The point here is that the higher amounts of D corresponds to the higher amount of the light-sensitive cells being spread within the tissue; therefore, more cells would be activated once an optical stimulation be induced, and consequently, a uniform simultaneous inactivation period within a significant number of cells of the tissue makes the fibrillation wavefront silent.

Furthermore, comparing two amounts of P, our study suggests that P = 0.4 (PL) results in more successful cases of defibrillation in comparison with their counterparts. For instance, DLPH results in 25% while DLPL results in 42.85% of successful cases. Examining other choices for P and D in future studies, will result in a better view for understanding the role of distribution of the light sensitive cells in the success/ failure of defibrillation procedure via optogenetic methods.

As the last part of this study, we induced a second optical stimulation at time step 15000 (i.e., 1.5 msec after the first stimulation) for the DLPL group to investigate the role of primary (former) stimulation. We observed that the percentage of successful cases after a quick stimulation is higher than when one single stimulation is applied later. Our results show a roughly 15% rise in the successful cases. The point at issue is that the time of stimulation could have an important role in the success of defibrillation.

## 5. Modeling Simplification Issues

There are two issues to be considered in this study. First, we utilized a simplified model for each cell, and second, our study was performed in a 2-D tissue model. Both simplifications were accepted to help us to reduce the cost of computations. Therefore, our results can be considered as preliminary results giuding us towards the main answers. As our future work, we will extend the proposed model to 3D. For example, in [22] it has been shown how to extend the fibrosis representation methods to 3-D [22]. However, using a more realistic model like TTM to represents the cells, is still under debate for us, because of computational costs it will cause for us.

## 6. Conclusions

In conclusion, we developed a low-order model for studying cardiac tissue. Because of the low order computations of the model, without common computational challenges, could be extended to a three-dimensional model. Our in-silico simulations suggest that a higher amount of relative area, entropy, and also transmurality may well result in more unsuccessful optogenetic defibrillations. However, we have not proved any meaningful relative between interface and successful defibrillation. Moreover, our simulations show that a higher amount of D, and also early induction time would be beneficial for the success of defibrillation procedure.

## List of Supplementary Materials

1. **Table S1** _ Average and interquartile amounts of the data set.

2. **Fibrillation Movies**_ Including 4 .avi files of sample fibrillations

3. **Defibrilation Movies**_ Including 5 .avi files of sample fibrillations

